# Three-dimensional models of the cervicovaginal epithelia to study host-microbiome interactions and sexually transmitted infections

**DOI:** 10.1101/2021.11.04.467382

**Authors:** Vonetta L. Edwards, Elias McComb, Jason P. Gleghorn, Larry Forney, Patrik M. Bavoil, Jacques Ravel

## Abstract

Two-dimensional (2D) cell culture systems have provided controlled, reproducible means to analyze host-pathogen interactions. Although inexpensive, straightforward, and requiring very short time commitment, these models recapitulate neither the functionality of multi-layered cell types nor the microbial diversity of an infected human. Animal models have commonly been used to recreate the complexity of human infections. However, extensive modifications are commonly required to recreate interactions that resemble those in the human reproductive tract microbiologically and physiologically. Three-dimensional (3D) cell culture models have emerged as alternative means of reproducing key elements of human infections at a fraction of the cost of animal models and on a scale that allows for replicative experiments to be readily performed. Here we describe a new 3D model that utilizes transwells with epithelial cells seeded apically and a basolateral extra cellular matrix (ECM)-like layer containing collagen and fibroblasts. In this system, basal feeding creates a liquid/air interface on the apical side. The model produced tissues with close morphologic and physiological resemblance to human cervical and vaginal epithelia, including observable levels of mucus produced by cervical cells. Infection by both *Chlamydia trachomatis* and *Neisseria gonorrhoeae* was demonstrated as well as the growth of bacterial species observed in the human vaginal microbiota, enabling controlled mechanistic analyses of the interactions between host cells, vaginal microbiota and STI pathogens. Future experiments may include immune cells to mimic more closely the genital environment. Finally, the modular set up of the model makes it fully applicable to the analysis of non-genital host-microbiome-pathogen interactions.

**IMPORTANCE:** Infected sites in humans are a complex mix of host and microbial cell types interacting with each other to perform specific and necessary functions. The ability to understand the mechanism(s) that facilitate these interactions, and interactions with external factors is paramount to being able to develop preventative therapies. Models that attempt to faithfully replicate the complexity of these interactions are time intensive, costly, and not conducive to high throughput analysis. Two-dimensional (2D) models that have been used as a platform to understand these interactions, while cost effective, are generally limiting in experimental flexibility and structural/physiological relevance. Our three-dimensional (3D) models of the cervicovaginal epithelium can facilitate analysis of interactions between the host epithelium, sexually transmitted pathogens and bacteria present in the vaginal microbiota. Due to the modular design, additional cell types and environmental modulators can be introduced to the system to provide added complexity, approaching conditions in the infected human host.

---

Eukaryotic cell culture systems have been a staple of host-pathogenesis research for decades as they provide the means to model *in vivo* interactions in a controlled and reproducible *in vitro* environment. The mainstay of this approach is a flat surface 2-dimensional (2D) model where cells are grown as a monolayer on a solid impervious surface, usually plastic or glass treated with polymers that enhance cell adhesion. This method is inexpensive, accommodates many adherent cell types and imaging of the cells is relatively straightforward [1-3]. While 2D models have provided a wealth of information on host-pathogen interactions, they do not faithfully reproduce the physiological complexity of these interactions as they occur within or on the host organism. Notably, 2D cell culture systems may not be representative of *in vivo* cell morphology, lack true cellular junctional complexes and fail to account for the effect of differing cell types usually found within the environmental milieu [3, 4]. Experimentally, the design of the 2D systems is also limiting, as it does not allow the introduction of an air interface or the incorporation of extracellular matrices (ECM) that produce needed signaling and crosstalk molecules. This means that many of the predictions derived from 2D cell culture models do not hold true when applied to *in vivo* situations, as seen in cervical cancer models [5], and other pathogen-host models [6, 7].

To overcome these obstacles, models for multiple diseases and conditions have been developed in animals. These afford the ability to follow a progressing infection in a complex environment that can replicate many properties of the human host, e.g., local physiology and host response, but falls short on many others e.g., the structural and polymicrobial environments. Additional manipulations and modifications are also often required to maximize susceptibility to human-specific infectious agents [8-13]. The use of animal models for STI research is further complicated by the need to use animal-adapted pathogens strains, as is the case with *C. trachomatis*, or alternate species that have coevolved with their host, as is the case with *Chlamydia caviae* and *Chlamydia muridarium* [14-17]. Lastly, animal models are often expensive to develop and maintain. This high cost may limit the number of replicate experiments and thus exhaustive investigations are not usually undertaken.

Three-dimensional (3D) cell culture models provide a practical, cost-effective alternative to animal models while also greatly improving the modelling value of 2D culture systems. 3D models can capture many aspects of the native *in vivo* physiology including cell morphology, organization, and communication that cannot be replicated in typical 2D models. This includes, but is not limited to, the ability to replicate complex tissue interactions, create and maintain intercellular interactions including junctional complexes, facilitate differentiation and polarization, mimic cellular behavior and integrate the site-specific microbial environment [4, 18-26]. Over the past decade various 3D cell culture reproductive tract model systems have been developed. These range from hydrogels [25, 27], and self-assembled organoids [3, 28], to microfluidics organ-on-a-chip models [23]. Hydrogels are usually placed on a scaffold and cells can be grown within or on top of the hydrogel, with 7 to 21 days necessary for full differentiation. These models have been used to analyze bacterial growth patterns, as well as targeted aspects of pathogenicity for multiple pathogens including ZIKA, HSV, *Chlamydia* spp., *Neisseria gonorrhoeae* and HIV [22, 29-34]. Similarly, self-assembled organoids can be grown either on a scaffold (i.e., collagen-coated beads) or scaffold free where cells are placed in suspension and self-aggregate to form a more complex structure. Organoid-based models have been used for mechanistic studies of bacterial pathogenesis [21, 28, 35-38]. 3D models generally closely mimic infections of multiple pathogens [22, 27, 30, 34, 36, 39] as well as environmental parameters [8, 29, 40] as they occur *in vivo*.

Whereas advances in hydrogel-based and organoid-based systems can recapitulate the 3D environment and multicellular nature needed to mimic aspects of the *in vivo* context, to an extent reproducibility is difficult owing to their stochastic cellular organization and/or time needed to establish the model. Organ on-a-chip models can overcome some of these limitations. Microfluidic modules that integrate parameters such as flow, mechanical stress, and the introduction of multiple environmental cues in any orientation around the cell(s) of interest can be developed [41-44]. This allows for organ-like systems that can be functionally maintained for extended periods of time allowing for more in-depth analysis. [20, 23]. However, the high cost of set up and maintenance of some of these models may not be feasible for many laboratories interested in studying host-pathogens interactions of the female reproductive tract. Indeed, there have been limited efforts towards the development of organ-on-a-chip systems to model infections of the reproductive tract.

In this study, we developed and characterized a 3D transwell cell culture model characterized by morphologically and physiologically differentiated vaginal and cervical epithelial cells that support the growth of bacteria found in the vaginal milieu and enable infection by both *C. trachomatis* [45] and *N. gonorrhoeae*. The transwell polyester membrane provides scaffolding support for the epithelial cells while allowing close proximity to an ECM and fibroblast network. By using the non-cancerous, mucin producing cell line (A2EN) [46], the model recapitulates critical aspects of the *in vivo* environment where mucins play an important role [47, 48]. Relatively low cost and short set-up time required to establish the model, enables the testing of multiple replicates in parallel under multiple conditions in a semi-high throughput process. This model may serve as a primer for the future development of more elaborate 3D organ-on-a-chip model systems.

## RESULTS

### The epithelial 3D transwell model structurally resembles *in vivo* cervical and vaginal epithelium

Transwells, basally coated with collagen and in the presence of human fibroblasts, mimicking the *in vivo* basement membrane of the epithelium, were used to develop a 3D model of the female reproductive tract epithelia (Fig. 1). In these models, basal only feeding and air interface exposure afforded the establishment of vaginal and cervical epithelia that morphologically closely resemble the structure of these epithelia *in vivo* (Figs. 2 and 3). Polarization of epithelial cells over time is usually an indication of their stage of development. We initially tested multiple types of collagen and coating methods to determine optimal conditions. Epithelial barrier integrity was evaluated using TEER values [49] measured over culture as the epithelial tissue formed. Basal collagen coating and fibroblast embedding showed a gradual increase in TEER values with peaks of approximately 600 ohms/cm^2^ on day 6 for A2EN cervical epithelial cells (Fig. 2A) and 1000 ohms/cm^2^ on day 8 for VK2 vaginal epithelial cells (Fig. 3A). Other methods of collagen coating and fibroblast embedding, including apical coating (Figs. 2A and 3A), as well as basal or apical coating with embedded fibroblasts were tested (data not shown). Embedding of fibroblasts was detrimental to the integrity of the collagen layer and caused delamination from the transwell membrane.

**FIG 1.**
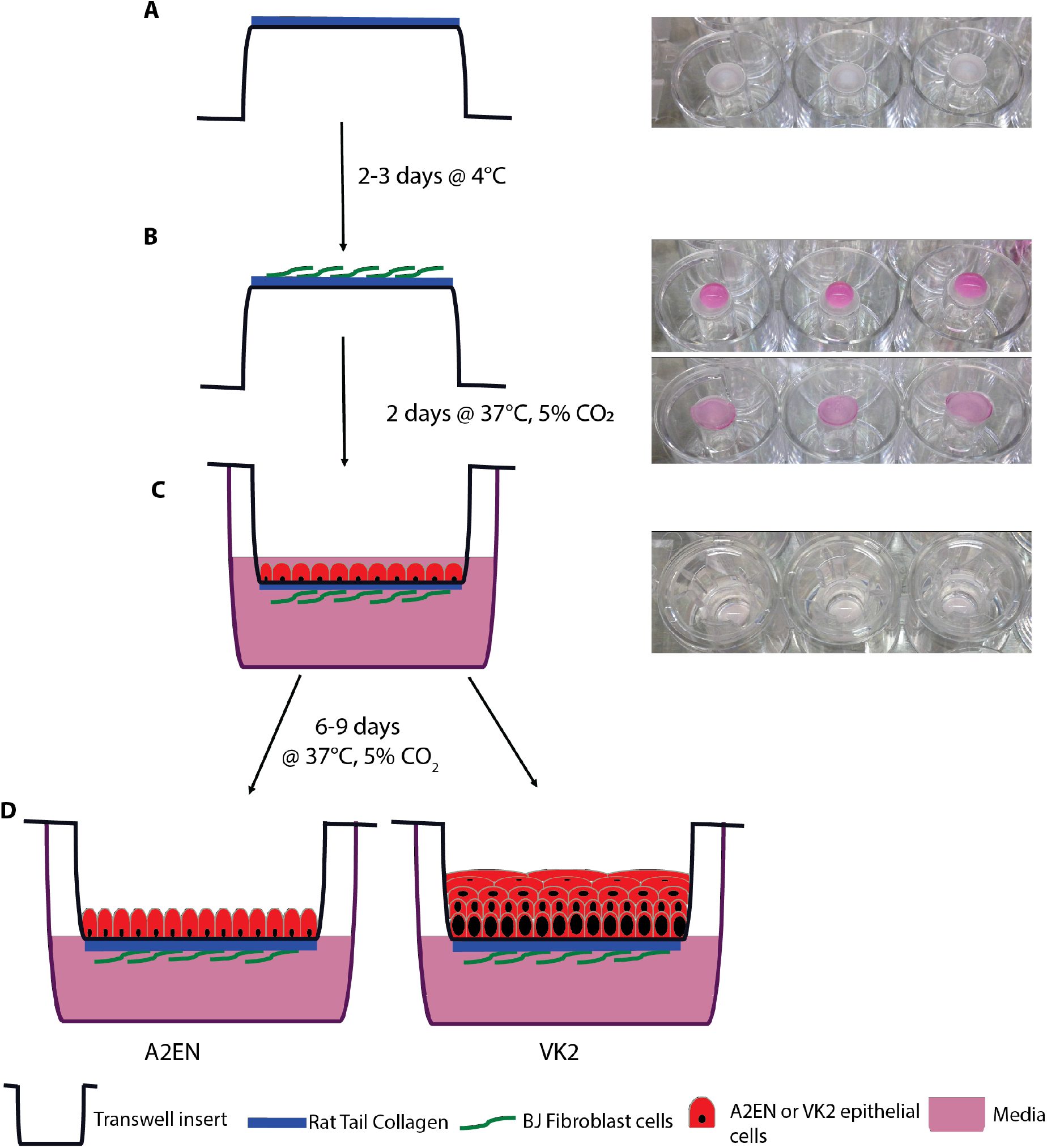
Model setup. 70μl of collagen was added to the basal portion of the inverted transwell (A) and stored at 4°C. BJ’s were added to the basal membrane 2-3 days later at 3×10^4^ in a volume of 80-100μl and incubated at 37°C, 5%CO_2_(B). Epithelial cells (A2EN or VK2) were apically added at 1×10^5^ in a volume of 50-200μl (C). After 6-9 days incubation at 37°C, 5%CO_2_ cells were ready to be used in experiments.

**FIG 2.**
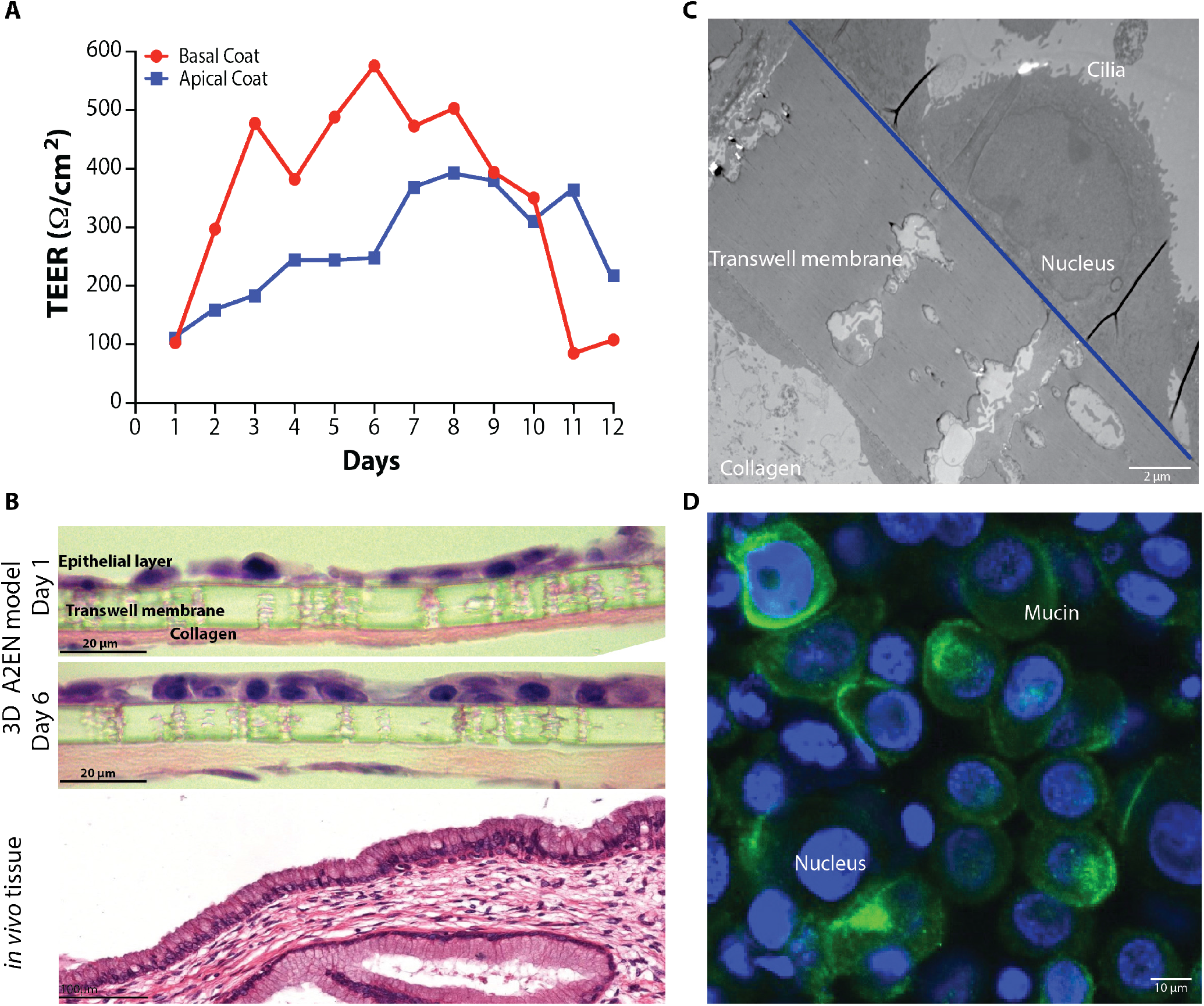
Characterization of the 3D cervical epithelium model (A2EN). Transepithelial resistance values over the course of A2EN epithelial cell transwell 3D model set up (A). Histology (H&E) imaging (B) and electron microscopy (TEM) imaging (C) of the epithelial cells of the model 6 days post set up. Confocal imaging of mucin gel formation (MUC-5B) on the model 6 days post set up (D).

**FIG 3.**
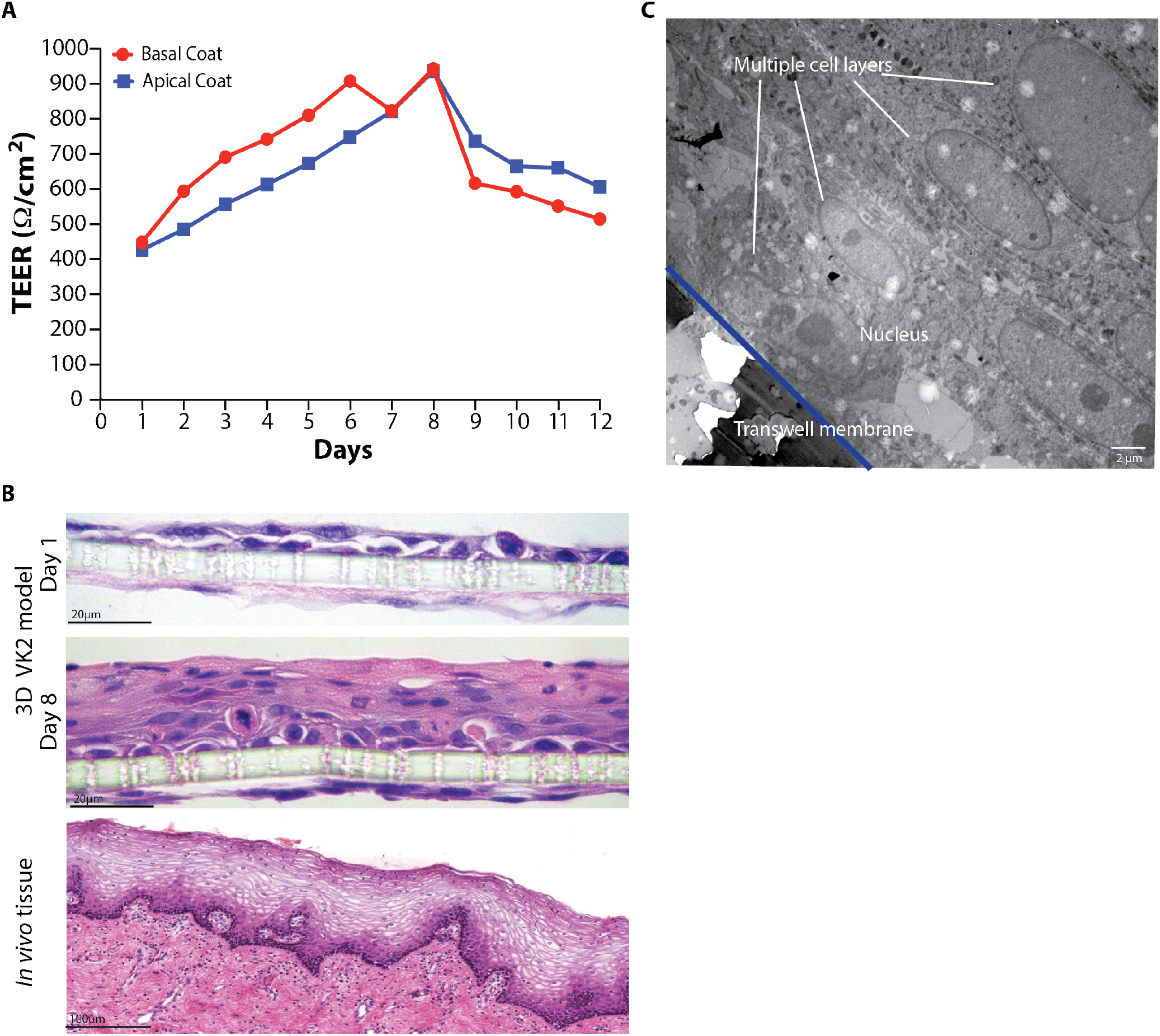
Characterization of the 3D vaginal epithelium model (VK2). Transepithelial resistance values over the course of VK2 epithelial cell transwell 3D model set up (A). Histology (H&E) imaging (B) and electron microscopy (TEM) imaging (C) of the epithelial cells of the model 8 days post set up.

Histology and electron microscopy were used to evaluate the structural and morphological features of the two epithelial cells models. Hematoxylin and eosin (H&E) staining confirmed the increased cell density and polarization of A2EN cervical epithelial cells (Fig. 2B) on day 6 as compared to day 1. Morphologically, the epithelial structure was similar to that observed in histological images of cervical tissue which comprises a compact single layer of epithelial cells (Fig. 2B). Transmission electron microscopy (TEM) further confirmed these observations. The A2EN cervical epithelial cells form a monolayer of cells in tight contact with each other. An intact nucleus and cilia on the surface of the cell were also observed (Fig. 2C).

An important feature of cervical cells is their ability to produce mucus [50, 51]. A2EN cervical epithelial cells were selected for their demonstrated ability to produce mucus in a 2D model; a unique and important feature of this cell line [46]. Immunostaining for mucin 5B, a major protein component of mucus, shows that mucus is produced over a significant portion of the apical surface of the A2EN cervical epithelium (Fig. 2D), thus recapitulating a critical functional property of the cervical epithelium [51].

Histology and electron microscopy imaging of the VK2 vaginal epithelial model revealed a pronounced stratification at day 8 as compared to day 1 with multiple layers (up to 7) of cells observed (Fig. 3B and 3C). Further, maturation of VK2 epithelial cells was observed, with more mature cells in the upper layers and more immature cells in the lower layers. This organization mimics an integral feature of the vaginal epithelium, as glycogen, a key metabolite supporting the growth of the vaginal microbiota, accumulates in mature epithelial cells [52].

### 3D cervical A2EN cells are infected by *Chlamydia trachomatis* and *Neisseria gonorrhoeae*

The ability of *C. trachomatis* (Ct) serovar L2 to infect cervical A2EN cells was assessed in both a conventional 2D model (Fig. 4A) of cells grown on coverslips and in the 3D transwell model described herein (Fig. 4B). While both models facilitated relatively robust infectivity (Fig. 4C), the A2EN cervical epithelium 3D model accommodated higher infection (71%) compared to the 2D model (57%) (p-value 0.019). Both models were both infected with 2×10^5^ *C. trachomatis* elementary bodies representing a MOI of 2 and 1 for the 2D and 3D models respectively. Since the MOI was lower for the A2EN cervical epithelium 3D model, it demonstrated that a more efficient infection can be achieved in that model. The VK2 vaginal epithelium 3D model was also successfully infected with *C. trachomatis* (data not shown); however, as expected the level of infectivity was low (25%) since *C. trachomatis* predominantly infects cervical epithelial cells. TEM confirmed the infection, visualizing inclusions containing *C. trachomatis* at various developmental stages (Fig. 4D), with both elementary bodies (EBs) (infectious particles) and reticulate bodies (RBs) (metabolic/replicating particles) observed. While A2EN cervical epithelial cells are not robust producers of cytokines [46], we investigated the profiles of some common cytokines and found appreciable levels of IL-6, IL-8, IP10 and RANTES (Fig. 4E). These results are similar to those observed in Buckner et al [46], where perceptible levels of IL-6, IL-8, IP10 and RANTES were detected. We observed that the cytokine response of the model in the presence or absence of a chlamydial infection was similar to that previously observed in this and other human and mouse cell lines [53-56]. These results indicate that the A2EN cervical epithelium 3D model could serve as a suitable platform for studies of chlamydial infection.

**FIG 4.**
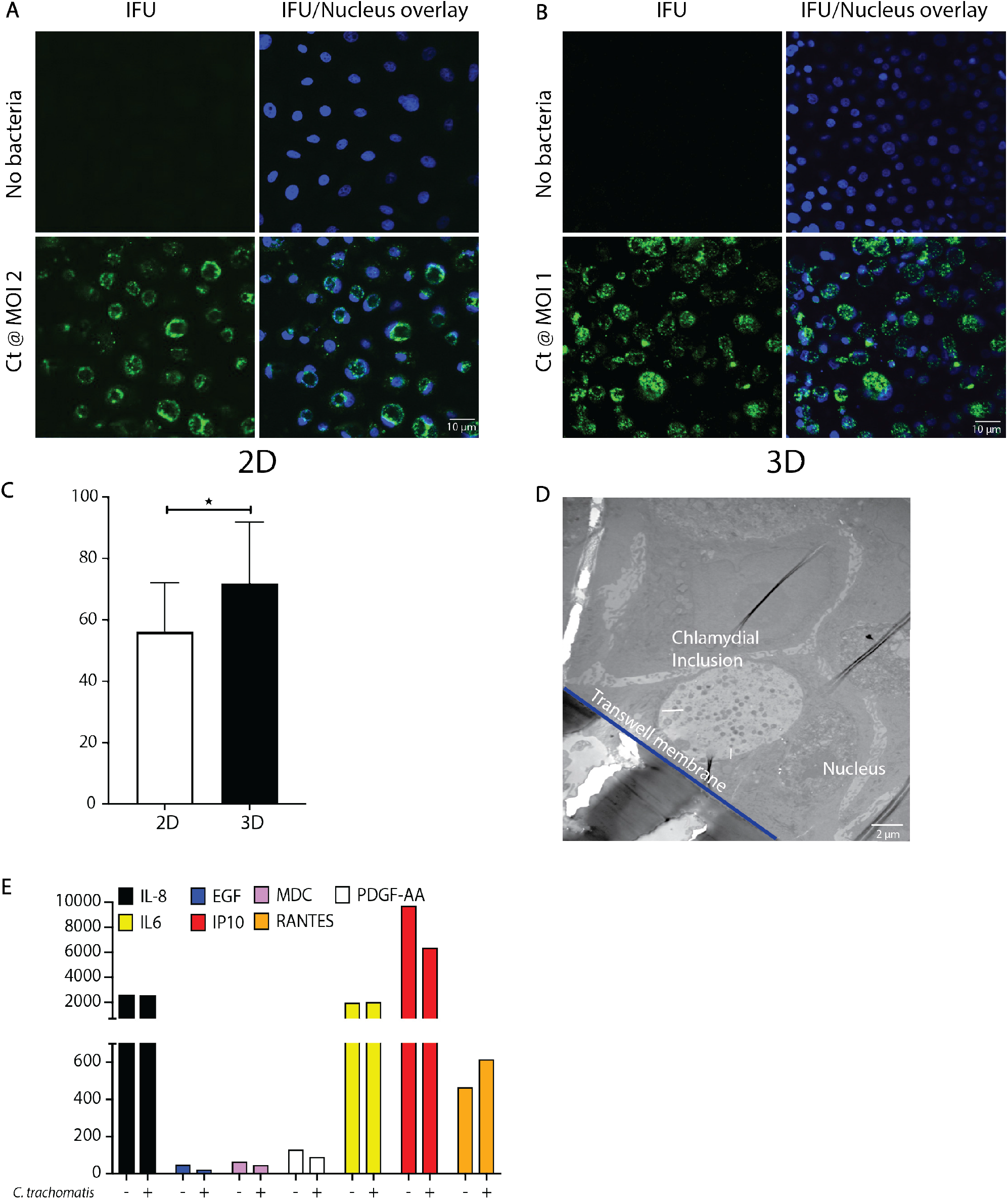
Infection of the 3D cervical model (A2EN) by *C. trachomatis*. Analysis of chlamydial infectivity on the conventional 2D (coverslip) model by fluorescent imaging (A) compared to the 3D (transwell) model (B) and resultant enumeration of infected cells (C). TEM image of infected cells on the 3D model (D). Cytokine profile of uninfected as compared to infected 3D cervical cells (E).

Another common sexually transmitted pathogen is *N. gonorrhoeae* [57], with anecdotal evidence suggesting that *N. gonorrhoeae* infection might lead to an increased risk of *C. trachomatis* infection [58, 59]. Utilizing wildtype and mutants of a common *N. gonorrhoeae* laboratory-adapted strain FA1090, we showed that transmigration of *N. gonorrhoeae* takes place within 6 hours in the 3D cervical A2EN model. This is similar to the transmigration period observed with a HEC-1-B 3D cell model (Fig. 5A), a cell line commonly used to analyze *N. gonorrhoeae* infections [60-63]. TEM imaging shows *N. gonorrhoeae* attached to the surface of the A2EN cells (Fig. 5B), which is the first step in the pathogenic cycle. These results suggest that the 3D A2EN cervical epithelium model can also support investigations of *N. gonorrhoeae* pathogenesis.

**FIG 5.**
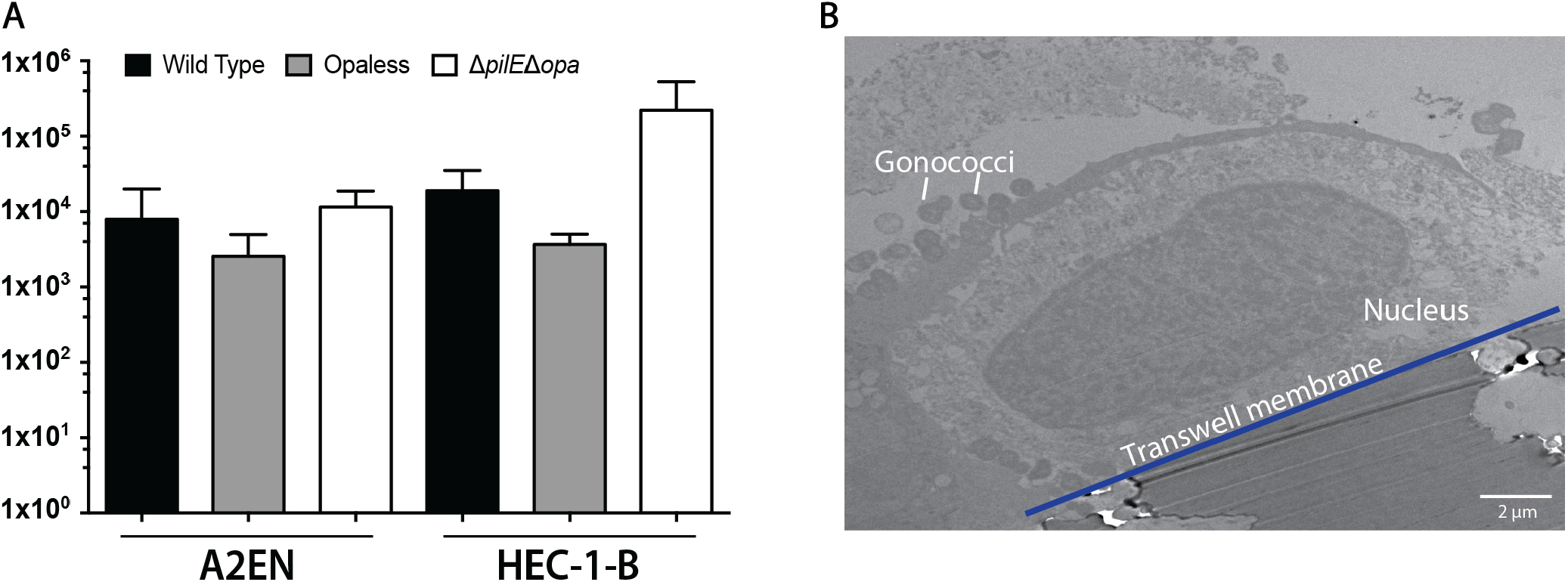
Infection of the 3D cervical cells (A2EN) by *N. gonorrhoeae* (Gc). Transmigration of Gc across the cervical epithelium model is similar to that obtained with a commonly used cell line (HEC-1-B) (A). TEM image of Gc attached to 3D cervical epithelial cells (B).

### The 3D vaginal model can sustain the growth of vaginal bacteria

The vaginal microbiota plays a key role in the cervicovaginal microenvironment [64]. We developed conditions that afford the growth of *Lactobacillus crispatus* and *Gardnerella vaginalis* on the 3D vaginal epithelium model. These two species are prominent members of vaginal bacterial communities that are found in optimal and non-optimal conditions, respectively [65]. These bacteria were used to inoculate on the 3D vaginal epithelium model and shown to grow for at least 48h under anaerobic conditions. Growth was first demonstrated by measuring the pH of culture medium in the apical compartment after 48h of growth. Media containing *L. crispatus* had a significantly lower pH of ∼4.2 as compared to *G. vaginalis* (pH 6.0) (Fig. 6A), As expected, *L. crispatus* acidified the microenvironment, while *G. vaginalis* did not. *In vivo* acidification is driven by the production of lactic acid by *L. crispatus*, typified with a higher proportion of D(-) lactate as compared to L(+) lactate [66-68]. A concentration of 7.41mM D(-) lactic acid was observed after 48h of growth with *L. crispatus* compared to 2.42 mM and 2.04mM with *G. vaginalis* or a no bacteria control, respectively (Fig. 6B). This finding demonstrates that *L. crispatus* is metabolically active and growing on the 3D vaginal epithelium model. Further microscopic analyses using both TEM (Fig. 6C i, ii, iii) and FISH (Fig. 6C iv, v, vi) showed the presence of live *L. crispatus* (Fig. 6C ii, v) and *G. vaginalis* (Fig. 6C iii, vi) on the model under anaerobic conditions after 48h growth. It is important to note that the model was gently rinsed with PBS before fixation thus any non-adherent bacteria were removed, only bacteria attached to the epithelial surface or embedded in the mucin matrix were imaged. TEM afforded visualizing the physical localization of the bacteria in close proximity to the epithelial layer, while FISH staining confirmed the robust growth of the bacteria on the model. Viability staining (Fig. 6C vii, viii, ix) after 48h of bacterial growth indicated that vaginal epithelial cells remained viable (green staining on Fig. 6C), in contrast to a control comprising of epithelial cells exposed to 1% saponin which predominantly stain red and indicate dead cells (Fig. 6C x).

**FIG 6.**
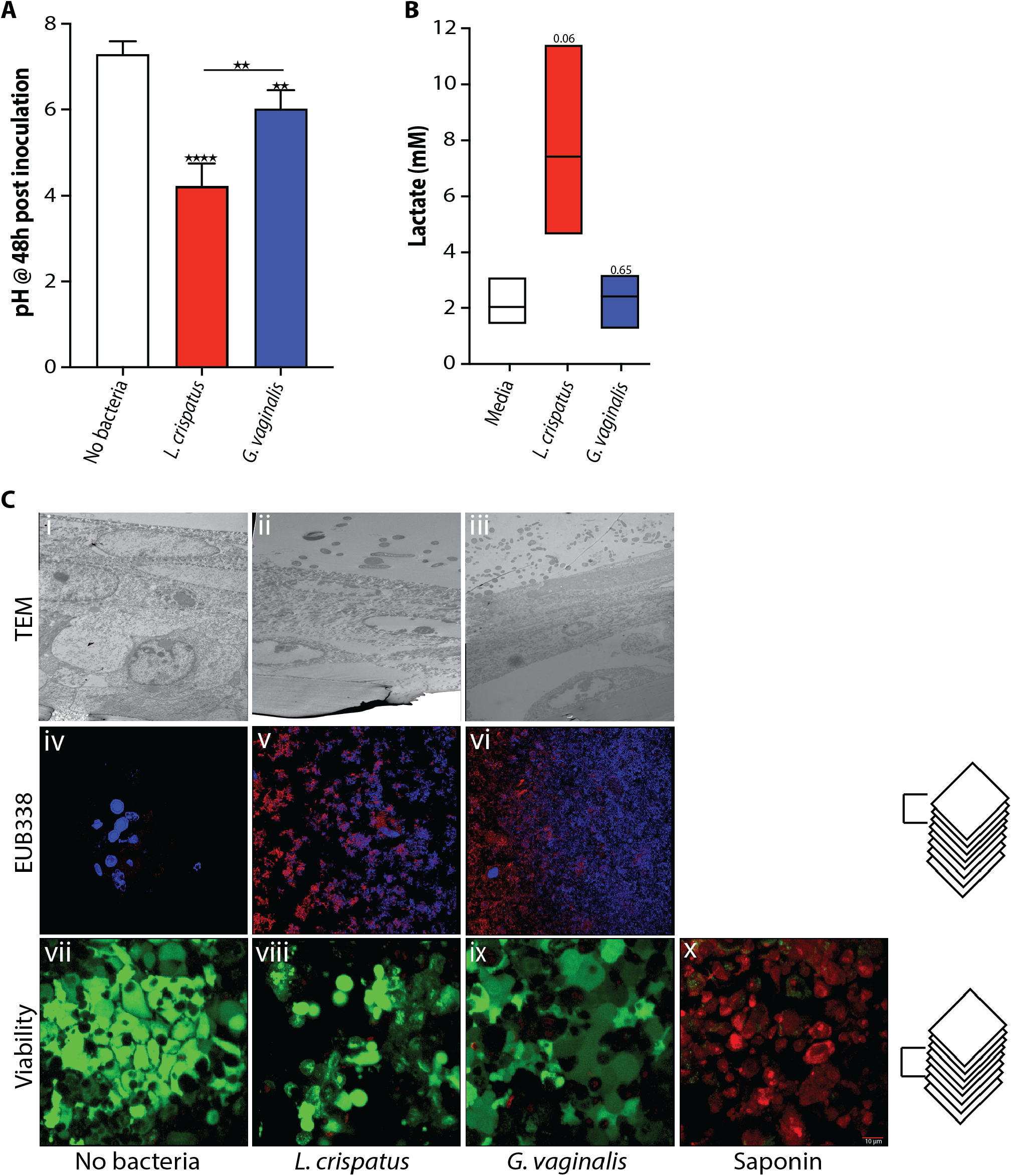
The 3D vaginal (VK2) epithelium model supports the growth of vaginal bacteria. pH (A) and D(-) lactate concentrations (B) of apical media after 48h of anaerobic bacterial growth on 3D VK2 cells. TEM, FISH and viability images of bacteria and host cells after 48h of growth (C).

## DISCUSSION

2D cell culture models have been extensively used to study host-STI pathogen interactions. However, these models lack complexity and do not accurately mimic many of the physiological interactions that occur in host organisms, limiting interpretation and translation to complex human physiology. More specifically, 2D culture models are unable to recapitulate the native microenvironment including multicellularity, the composition of extracellular matrices (ECM), or various physicochemical properties and spatiotemporal molecular gradients [4, 69]. As such, many of the predictions derived using these 2D cell culture models often do not hold true when applied to conditions *in vivo*, as seen in cervical cancer models [5] and other pathogen-host models [6, 70]. We have advanced these models by developing 3D organotypic models of cervical and vaginal epithelia that include an interstitial compartment of collagen and associated fibroblasts. 3D epithelial models provide enhanced morphological and physiological cellular structures that can include inter-cellular interactions (i.e., junctional complexes), complex tissue interactions, differentiated and polarized epithelial structures, which taken together better mimic *in vivo* cellular behavior [4, 18-23, 25, 28]. The multi-layered structure of these models affords increases in complexity and experimental flexibility, such as the potential addition of different cell types, or even immune cells. 3D cell culture models partly fill a gap between the cost effectiveness of 2D cell culture and the complexity and high cost of organoids, organ-on-a-chip systems or animal models [8, 9, 11-13]. Animal models can be of limited use to study host-STI pathogen interactions because they are often lacking anatomical similarity to the human vaginal epithelium. For example, the lower reproductive tract of the mouse, an animal model commonly used in STI research, comprises of a keratinized stratified epithelium, while that of human is not keratinized. The 3D organotypic model we have developed is ideally suited for studies on the pathogenesis of STIs as it replicates many features of human cervicovaginal epithelia without the complexity, experiment-to-experiment variability and/or cost of organoids and animals. This proposed model will ultimately provide a way to study how the cervicovaginal microbiota interact with the host, and how these interactions increase or reduce the risk of infections by sexually transmitted pathogens.

We have shown that the model supports infection by *C. trachomatis* and *N. gonorrhoeae*, two of the most prevalent infections worldwide. *C. trachomatis* is an obligate human pathogen that requires host internalization for propagation while *N. gonorrhoeae* can replicate both outside and inside of epithelial cells. The 3D cervical model was able to reproduce characteristic features of infection by both pathogens. One can envision using these models to study co-infections or the role of a primary infection by *C. trachomatis* in susceptibility to infection by *N. gonorrhoeae*, or vice versa. Other potential co-infections, including with HSV, HPV or HIV could also be investigated. The model can be enhanced further by adding more complex structures such as endothelial and/or immune cells to the basal compartment.

Importantly, the ability to grow vaginal bacteria on the 3D models of the vaginal and cervical epithelia is a critical first step toward modeling the *in vivo* complex microenvironment that includes a functional microbiota. Little is known about how the vaginal microbiota contributes to modulating the risk to STIs. Previous studies have postulated that indole-producing bacterial species such as *Prevotella, Petpostreptococcus* or *Peptinophilus* spp. can facilitate *C. trachomatis* replication [71-73], since *C. trachomatis* can use indole to synthesize tryptophan, an essential amino acid that genital *C. trachomatis* strains are incapable of producing. Tryptophan is present in the host extracellular and cytoplasmic compartments but can be depleted through the action of interferon (IFN)-γ which induces tryptophan catabolism by indoleamine-2,3-dioxygenase I (IDO) [74]. Mechanisms such as this have been difficult to study for at least three reasons: (1) it is unethical to perform many of these experiments in humans; (2) there are no cellular or biomimetic models of the cervicovaginal environment that include the microbiota; and (3) key features of the cervicovaginal space such as the dominance of *Lactobacillus* spp. and a low environmental pH (<4.5) are not found in other mammals that might otherwise be candidate animal models [75-77]. The 3D models developed in this study represent the first steps toward more advanced models that include complex microbiota. This component is critical, as the cervicovaginal microbiota exists in a mutualistic relationship with the cervicovaginal epithelium and is believed to play an important role in the risk to STIs. The microbiota is thought to constitute the first line of defense against STIs, but the mechanism(s) by which it exerts its protective effect(s) is/are unknown. Access to a model that reproduces the physiology and microbiology of the cervicovaginal space is thus critical. We have previously shown that an optimal microbiota dominated by *Lactobacillus* species, such as *L. crispatus*, produces copious amounts of lactic acid and a concomitant low environmental pH (<4.5). Lactic acid does not directly affect *C. trachomatis* bacteria but acts on the epithelium by decreasing epithelial cell proliferation, thus significantly inhibiting the infection process [45]. On the other hand, microbiota compositions associated with an increased risk to STIs tend to be similar to those observed in association with bacterial vaginosis (BV). BV is a condition that is generally defined by a high pH (>4.5), a microbiota characterized by the absence of *Lactobacillus* spp. and the presence of an array of strict and facultative anaerobes such as *G. vaginalis, Atopobium vaginae*, and *Prevotella* spp. The mechanisms by which a STI-permissive microbiota increase the risk to infection remains poorly understood. Based on our previous research we posit that a non-permissive indigenous microbiota interacts with the cervicovaginal epithelium to establish a homeostatic state that blocks STI and/or reduces disease severity. Conversely, we propose that a permissive microbiota disrupts host epithelial cell homeostasis, thereby allowing STI to progress. Establishing reconstituted STI-permissive or non-permissive microbiota on an advanced 3D epithelial models will go a long way toward testing these hypotheses and improving our knowledge of the pathogenesis of STIs.

The 3D models developed in this study uses relatively inexpensive materials compared to organoids or organ-on-a-chip systems. These low-cost models afford performing replicate experiments in a semi high-throughput setup. In addition, this 3D model allows performing different analyses from one or replicate transwells, including resistance readings, measurements of pH, metabolite concentrations (i.e., lactate), cytokine concentrations, bacterial enumeration, and imaging (fluorescence, TEM) or omic analysis (DNA/RNA sequencing, proteomics, among others). Lastly, while we developed this system with A2EN cervical and VK2 vaginal cell lines, there is no barrier to using different cell lines more appropriate to the research questions at stake, or even from different organ systems or tissues.

## MATERIALS AND METHODS

Abbreviations and all catalog numbers are listed in the supplemental materials.

### Cell Culture Model: collagen coating

Transwell inserts (Corning #3472) were removed from the 24-well plate using glass pipettes or tweezers and placed in an inverted orientation into 12-well plates. To form the collagen coating, all solutions were chilled and placed on ice. 200μl 5X RPMI (1:1 mixture of 10X RPMI and tissue culture (TC) water) (Sigma #R1145) and 25μl 1M NaOH (Sigma #S5881) were combined and vortexed thoroughly for 10 secs. Rat tail collagen (800μl; Corning #354236) was added with gentle pipetting to avoid introducing excessive bubbles, and the pH of the mixture was tested. Additional NaOH or RPMI was added in 1μl or 10μl increments if needed to attain a pH of 6.5 (the final mixture should have a salmon pink color). A total of 70μl of the collagen mixture was added to the basal surface of each transwell insert, the plate was covered ensuring no contact with the collagen surface and the collagen allowed to gel in a Biosafety Level 2 (BSL2) hood at room temperature for ∼30 min. (Fig. 1A). Using clean glass pipettes or tweezers the inserts were returned to the 24-well plate in the standard orientation and left under the hood for an additional 3h before transfer to 4°C for 48-72h.

### Cell Culture Model: addition of fibroblasts

After 48-72h, the transwells were inverted into a 12-well plate using glass pipettes or tweezers. BJ fibroblast cells (ATCC #CRL 2522) at 70-90% confluency after growth in BJ complete medium (DMEM media (Cellgro #15-013-CV) supplemented with 10% FBS (Sigma #F4135)) were trypsinized using 1ml of 0.25% Trypsin (Gibco #25200-056) and cell number determined using the Countess Automated Cell Counter. A total of 3×10^4^ cells in a volume of 75-80μl were added to the basal surface of the transwells on top of the collagen (Fig. 1B). The dish was covered and placed in a 37°C, 5% CO_2_ incubator for 6h. Inserts were then transferred to the 24-well plate in the standard orientation and BJ complete medium added (200μl to the apical compartment, 500μl to the basal compartment). The transwells were then returned to the incubator for an additional ∼42h.

### Cell Culture Model: addition of epithelial cells

Either A2EN cervical epithelial cells (kindly provided by Dr. Allison Quayle [46]) or VK2/E6/E7 vaginal epithelial cells (ATCC #CRL 2616) were used to make cervical or vaginal models respectively. A2EN cervical cells were grown in A2EN complete medium (EpiLife media (Gibco #MEPICFPRF) with 100X EDGS supplement (Gibco #S-012-5) and 100X L-glutamine (Lonza #17-605E)) while VK2/E6/E7 vaginal cells (ATCC #CRL 2616) grown in VK2 complete medium (Keratinoctye-SFM (with BPE and EGF) (Gibco #10725-018) supplemented with 0.4M calcium chloride (Amresco #E506) and 100X L-glutamine (Lonza #17-605E)). Cells were grown until 70-90% confluent then trypsinized using 1ml of 0.25% Trypsin. The number of cells was determined using the Countess Automated Cell Counter. BJ complete medium was removed from the transwells which were then gently rinsed with 500μl PBS. To seed the epithelial cell layer, using A2EN or VK2 complete medium (cervical and vaginal model respectively) a total of 1×10^5^ epithelial cells in 200 μl of media was added to the apical compartment and 500 μl of media was added to the basal compartment and the plate returned to the incubator. After 48h the apical and basal media were removed by vacuum aspiration and fresh medium added to the basal compartment only, to create an epithelial-air interface. Fresh medium (500μl) was added to the basal compartment every other day. Following culture A2EN: 6 days and VK2: 8 days epithelial cells were polarized and no medium could be observed entering the apical compartment from the basal compartment, indicating stable epithelial barrier formation.

### *Chlamydia trachomatis* infection, microscopy imaging and cytokine analysis

*C. trachomatis* serovar L2 (strain LGV/434/Bu) was propagated in HeLa monolayers as previously described in Tan *et al*. [78]. Briefly, serovar L2 was cultivated in 100mm^2^ tissue culture dishes containing HeLa cells grown at 37°C, 5% CO_2_. Monolayers were gently rocked for 2h, fresh medium was added, and the infection was allowed to progress for 48h. Lysates were harvested, and inclusion-forming units (IFUs) calculated and stored in sucrose phosphate glutamate (SPG) [78] at -80°C. Seeds were used directly from -80°C stocks. *C. trachomatis* was inoculated at a multiplicity of infection (MOI) of 1 or 2.

The 3D model was inoculated with 50μl *C. trachomatis* in the apical compartment and rocked for 2h at room temperature. The *C. trachomatis* suspension was removed by pipetting, the cells were rinsed with 500μl PBS, fresh medium (500μl) added basally, and the model was incubated for an additional 46h at 37°C, 5% CO_2_.

Following infection, the transwells were prepped for imaging as described in the fluorescence staining section and images were obtained using a Zeiss Duo 5 confocal microscope and 3 consecutive Z stack slices were compressed to create images for confocal analysis. For electronic microscopy, the transwells were placed in glutaraldehyde fixative for processing and imaged on the Tecnai T12 Transmission Electron microscope. For comparative purposes A2EN cells were grown on coverslips (2D) for 2 days and then infected with *C. trachomatis* at MOI 2 [79]. Infection and staining were performed as described below with images obtained using a Zeiss Axio Imager Z1 (Zeiss). Infected cells were manually identified using the ImageJ software (NIH).

For cytokine analysis, medium was removed from the basal compartment and stored at -80°C. Seven cytokines: EGF, IL-6, IL-8, IP10, MDC, PDGF-AA and RANTES were analyzed using a Luminex Multianalyte assay at the UMB Cytokine Core Laboratory.

### *Neisseria gonorrhoeae* infection and analysis

3D models containing 2×10^5^ A2EN cells was exposed apically to 100μl of *N. gonorrhoeae* FA1090 wildtype or *N. gonorrhoeae* Opaless (all *opa* genes deleted) or *N. gonorrhoeae* Δ*pilE*Δ*opa* (*pilE* and all *opa* genes deleted) at MOI of 10 for 6h at 37°C, 5% CO_2_. The basal medium (500μl) was then removed and dilutions were plated on GCK agar plates to determine the number of *N. gonorrhoeae* bacteria that transmigrated within the 6h incubation period. For comparison 2×10^5^ HEC-1-B endometrial cells utilizing the same 3D set-up were exposed in parallel and the transmigrated *N. gonorrhoeae* bacteria were quantified.

### Bacterial growth (*L. crispatus, G. vaginalis*) and microscopy imaging

The optical densities (OD) of bacterial cultures grown overnight in their respective media (*L. crispatus* in NYCIII and *G. vaginalis* in TSB+5% horse serum) were determined using an OD to colony forming units (CFU) conversion of 1 OD represents 1×10^9^ CFU. A volume corresponding to 2×10^8^ CFUs was added to the experiment culture medium (a 2:1 mixture of complete cell culture medium: bacteria culture medium) to produce a final volume of 1ml. A 10-fold dilution was then performed using experiment culture medium and 100μl was added to the apical compartment of the model (1×10^6^ CFU). Cells exposed to bacteria or medium only (no bacteria control) were incubated for 48h under anaerobic conditions in a 37°C incubator within a Coy chamber. Aliquots of media were removed from the apical compartment and the pH determined using an Apera Instruments PH8500 portable pH meter. Aliquots of 50μl were used to determine the D(-) and L(+) lactic acid concentrations using the Boehringer Mannheim/R-Biopharm D-Lactic acid/L-Lactic acid kit as per manufacturer’s instructions. Cells were gently rinsed with PBS and either fixed in 2.5% glutaraldehyde (TEM imaging) or 2% PFA (fluorescence imaging). Cells for TEM imaging were taken to the Electron Microscopy Core Imaging facility for further processing and imaging as described below. Cells for fluorescence *in situ* hybridization (FISH) imaging were processed as described below and imaged on a Zeiss Duo 5 confocal microscope and 5 consecutive Z stack slices were compressed to create images. Cells for histology and cell viability imaging were processed as described below and imaged on a Zeiss Duo 5 confocal microscope where 3 consecutive Z stack slices were compressed to create images (viability) or a Zeiss Primo Star (histology).

### Hematoxylin and Eosin (H&E) staining

The transwell membrane was excised from the support by rinsing the cell surface with PBS and cutting the perimeter of the membrane with a No. 11 blade on a scalpel. The membrane was then placed between two 32×25×3mm biopsy pads and secured in a histology cassette. The cassette was immersed in 10% formalin fixative solution for 24h and then processed by the UMB Pathology Histology Core using SOP NH306. Briefly, slides were placed in hematoxylin, rinsed with water, dipped in acid alcohol, rinsed with water, then sequentially placed in 80% ethanol, eosin, 95% ethanol twice, 100% ethanol twice and xylene thrice. Mounting media and a coverslip were then added. Resultant slices were imaged at 100X on the Zeiss Primo Star microscope (Zeiss).

### Fluorescence staining

Briefly, cells were rinsed once with 500μl of Dulbecco’s Phosphate Buffered Saline (PBS), fixed with 4% paraformaldehyde (PFA) for 30 min and permeabilized with 200μl 0.25% Triton X-100 in PBS for 10 min, followed by treatment with 300μl 0.1% Triton X-100 in PBS/Fish skin gelatin (FSG) (0.66%) for 20 min. The cells were then stained for chlamydial IFUs with 10μl of 5μg/ml of mouse anti-human chlamydia LPS (primary Ab) (US Biological, MA) in Triton X-100/PBS/FSG solution for 90 min. Secondary antibody staining was done by adding 2μl of 200μg/ml of goat anti-mouse Alexa Fluor 488 in Triton X-100/PBS/FSG solution and incubated 60 min in the dark. Host cells were stained with 2μl of 500μg/ml Hoechst in Triton X-100/PBS/FSG solution for 10 min in the dark. Chlamydial inclusions stained green while host cells nuclei stained blue. Cells were imaged using a Zeiss Duo 5 confocal microscope and 3 consecutive Z stack slices were compressed to create images for analysis.

### Transmission electron microscopy (TEM) staining

Cells were rinsed with PBS after removal of media and fixed in 500μl of 2% paraformaldehyde, 2.5% glutaraldehyde and 0.1 M PIPES buffer (pH 7.4) for at least 1 hour. Cells were then washed with 500μl of 0.1 M PIPES, quenched with 500μl of 50mM glycine in 0.1 M PIPES buffer (pH 7) for 15 minutes, washed and post-fixed in 200μl of 1% (w/v) osmium tetroxide and 0.75% ferrocyanide in 0.1M PIPES buffer at 4°C for 60 min. Following washing, transwell membranes were sliced off the holding cup, stained with 200μl of 1% (w/v) uranyl acetate in water for 60 min, dehydrated by passage through a graduated ethanol series and embedded in spurr’s resin (Electron Microscopy Sciences, PA) following the manufacturer’s recommendations. Resin blocks were trimmed perpendicular to the monolayer grown on the transwell membrane. Ultrathin sections ∼70nm thickness were cut on a Leica UC6 ultramicrotome (Leica Microsystems, Inc., Bannockburn, IL) and collected onto formva film coated SynapTek NOTCH-DOT grids (Electron Microscopy Sciences, Hatfield, PA) and examined in a Tecnai T12 transmission electron microscope (Thermo Fisher Scientific, formerly FEI. Co., Hillsboro, OR) operated at 80 keV. Digital images were acquired by using a bottom mount CCD camera and AMT600 software (Advanced Microscopy Techniques, Corp, Woburn, MA).

### Fluorescence *in situ* hybridization (FISH) staining

Cells were stained using a protocol modified from Meaburn et al. [80]. Briefly, cells were rinsed and fixed overnight at 4°C with 2% PFA then incubated in 200μl of 0.5% saponin/ 0.5% Triton X100/ PBS mixture for 40 min. This was followed by the addition of 200μl of 1N HCL for 20 min, 2X SCC for 10 min and 50% formamide/2X SCC for 30 min incubations. Cells were then incubated in 300μl of the hybridization mix containing the FISH probe EUB338-ATT0 for 10 min at 85°C then overnight in a humidity box at 37°C. Cells were washed with 500μl of multiple buffers (a) 50% formamide/2X SSC buffer at 45°C, (b) 1X SSC buffer at 45°C and (c) 0.05% Tween-20 in 4X SSC buffer at room temperature. Cells were then incubated with 300μl of Hoechst at 1:500 for 10 min and mounted for imaging. Cells were imaged on the Zeiss Duo 5 microscope (Zeiss) using the 63X objective with 488 and 546 filters.

### Viability staining

At 48h post-infection cells were incubated with 300μl of 4 μM Calcein-AM and 2 μM EthD-III from the Viability/Cytotoxicity Assay Kit for Animal Live and Dead cells (Biotium 30002-T) for 45 min at room temperature as per manufacturer’s recommendations. Cells were imaged at 40X using the 488nm and 543nm excitation wavelengths on the Zeiss Duo 5 microscope (Zeiss). A composite overlay of 3 Z stack slices were used to create a 3D image.

## ACKNOWLEDGEMENTS

The authors would like to thank Dr. Alison Quayle for kindly providing the A2EN cell line, Dr. Ruching Hsia and the staff of the University of Maryland-Baltimore Electron Microscopy Core for preparing cells for EM imaging. Dr. Joseph Mauban of the University of Maryland-Baltimore Confocal Microscope Core for advice and assistance on the confocal microscope. University of Maryland School of Medicine’s Center for Innovative Biomedical Resources, Histology Core and Cytokine Core – Baltimore, Maryland for providing advice, sectioning, and staining or running the samples for analysis.

## AUTHOR CONTRIBUTIONS

J.R., P.M.B. and V.L.E designed research; V.L.E. and E.M. performed research; V.L.E. and E.M. analyzed data; V.L.E., J.R., P.M.B., J.P.G., and L.J.F. wrote the paper; J.R., J.P.G. and P.M.B. obtained funding.

## Supplemental Materials

Transwell inserts (Corning #3472)

12 well plates (Corning #3513)

9” Glass pipettes (Corning #7095D)

Countess Automated Cell Counter (Invitrogen #C10227)

Mouse anti-human chlamydia LPS (US Biological #C4250-51F)

Goat anti-mouse Alexa Fluor 488 (Invitrogen #A-11029)

Rabbit anti-human MUC5B (Invitrogen #PA5-82342)

Goat anti-rabbit Alexa Fluor 488 (Invitrogen #A-11034)

Hoechst (Invitrogen #H3570)

FISH probe - EUB338-ATTO [5’-/565/GCT GCC TCC CGT AGG AGT-3’] (Invitrogen)

Histology fixative – 10% formalin solution (Sigma #HT501128)

D/L Lactic acid assay kit (R-Biopharm #11 112 821 035)

Rat tail collagen – 100mg (Corning #354236)

10X RPMI media (Sigma #R1145)

Sterile tissue culture water (Cellgro #25-055-CM)

1M NaOH – sterile filtered (Sigma #S5881)

0.25% Trypsin (Gibco #25200-056)

BJ human fibroblasts (ATCC #CRL 2522)

BJ complete media - DMEM media (Cellgro #15-013-CV) supplemented with 10% FBS (Sigma #F4135)

A2EN cervical epithelial cells (kindly provided by Dr. Allsion Quayle [46])

A2EN complete media - EpiLife media (Gibco #MEPICFPRF) with 100X EDGS supplement (Gibco #S-012-5) and 100X L glutamine (Lonza #17-605E)

VK2/E6/E7 human vaginal epithelial cells (ATCC #CRL 2616)

VK2 complete media - Keratinoctye-SFM (with BPE and EGF) (Gibco 310725-018) supplemented with Calcium chloride (Amresco #E506) and 100X L glutamine (Lonza #17-605E)

HeLa cervical epithelial cells (ATCC #CCL2)

HeLa complete media - Dulbecco’s modified Eagle’s medium (Corning #15-013-CV) supplemented with 10% FBS (Sigma #F4135)

HEC-1-B endometrial cells (ATCC #HTB-113)

HEC-1-B complete media – MEM alpha (1X) + GlutMaAX (GIBCO #32561-037) supplemented with 10% FBS (Sigma #F4135)

*Chlamydia trachomatis* serovar LGV II strain 434 (ATCC #VR 902B)

*Neisseria gonorrhoeae* strain FA1090 – wildtype and isogenic mutants (kindly provided by Dr. Alison Criss [81, 82]

*Lactobacillus crispatus* (ATCC #33197)

*Lactobacillus iners* (ATCC #55195)

*Gardnerella vaginalis* (ATCC #14018)

Difco GC medium base (BD #228920) with 1% Kelloggs supplement prepared as per White and Kellogg [83].

Bacteriological agar (Amresco #J637)

NYCIII medium: 10 g/L proteose peptone, 10 g/l beef extract, 5 g/l yeast extract, 5 g/L NaCl, 1.2 g/L MgSO_4_, 2 g/L MnSO_4_.H_2_O, 5.7 g/L K_2_HPO_4_, 20 g/L glucose, 10% FBS.

Tryptic Soy Broth (Fluka #T8907) supplemented with 5% Horse serum (GIBCO #26050-088)

